# dna-parser: a Python library written in Rust for fast encoding of DNA and RNA sequences

**DOI:** 10.64898/2026.01.20.700656

**Authors:** Matthieu Vilain, Stéphane Aris-Brosou

**Affiliations:** Department of Biology, University of Ottawa, 30 Marie Curie Pvt., Ottawa, K1N 6N5, Ontario, Canada; Department of Mathematics and Statistics, University of Ottawa, 30 Marie Curie Pvt., Ottawa, K1N 6N5, Ontario, Canada

**Keywords:** feature extraction, genomic sequences, Python library, encodings, machine learning

## Abstract

**Background:** The ever-growing amount of available biological data leads modern analysis to be performed on large datasets. Unfortunately, bioinformatics tools for preprocessing and analyzing data are not always designed to treat such large amounts of data efficiently. Notably, this is the case when encoding DNA and RNA sequences into numerical representations, also called descriptors, before passing them to machine learning models. Furthermore, current Python tools available for this preprocessing step are not well suited to be integrated into pipelines resulting in slow encoding speeds.

**Results:** We introduce `dna-parser`, a Python library written in Rust to encode DNA and RNA sequences into numerical features. The combination of Rust and Python allows to encode sequences rapidly and in parallel across multiple threads while maintaining compatibility with packages from the Python ecosystem. Moreover, this library implements many of the most widely used types of numerical feature schemes coming from bioinformaticss and natural language processing.

**Conclusion:** `dna-parser` is an easy to install Python library that offers many Python wheels for Linux (muslinux and manylinux), macOS, and Windows via pip (https://pypi.org/project/dna-parser/). The open source code is available on GitHub (https://github.com/Mvila035/dna_parser) along with the documentation (https://mvila035.github.io/dna_parser/documentation/).

## 1 Background

Since the mid-2000s, next-generation sequencing has led to a steep increase in the scales of datasets produced in biology (Muir et al. 2016). This revolution has led to the creation of publicly available databases such as the European Molecular Biology Laboratory Nucleotide Sequence Database (Kanz et al. 2005) or the Sequence Read Archive (SRA) (Leinonen et al. 2010), which grew significantly through time. The SRA in particular reached four petabases (4 × 10^15^ bases) around 2016 (Muir et al. 2016), twenty petabases in 2022, and is still growing exponentially (Edgar et al. 2022). These databases have enabled many discoveries and breakthroughs in biology in recent years despite their being challenging to analyze due to their sizes. For example, Edgar et al. (2022) discovered over 10^5^ new RNA viruses by analyzing data from public databases at scale. This data also fuels machine learning (ML) models for various applications **?**. For instance, by leveraging publicly available sequences of influenza, HIV-1, and SARS-CoV-2 viruses, Hie et al. (2021) accurately assessed which viral mutation can escape recognition from antibodies. A recent large language model also enabled the prediction of fitness changes of microorganisms based on changes in nucleotide sequences after training on 2.7 million prokaryotic and phages genomes (Nguyen et al. 2024). But, the most important recent breakthrough in biology involving ML and big data is arguably AlphaFold (Jumper et al. 2021), a model that uses over 2 billion protein sequences to accurately fold new proteins.

When building ML models that handle genomic sequences, the preprocessing steps are often of critical importance. Genomic sequences cannot be directly passed to models and usually have to be encoded into numerical features (a.k.a. descriptors). Many transformations exist and represent the sequences in different manners. For DNA and RNA sequences, some numerical representations are based on simple fixed mappings. It is the case for the real-number encoding where each base pair (bp) is represented as a real-number (*A* = *−*1.5, *T* = 1.5, C=0.5, *G* = *−*0.5) in a way that keeps the complementary nature of the bp (Chakravarthy et al. 2004; Kwan and Arniker 2009). This is also the case for the one-hot or Voss encoding (Voss 1992; Kwan and Arniker 2009; Yu et al. 2018) that maps each bp to a binary indicator to indicate its presence or absence. Other numerical features are based on physicochemical properties of the bp such as electron-ion interaction pseudopotential (EIIP) (Nair and Sreenadhan 2006), atomic number (Kwan and Arniker 2009) and Z-curve representation (Zhang and Zhang 2014). Additionally, techniques coming from the field of natural language processing like term frequency-inverse document frequency (TF-IDF)(Salton and Buckley 1988), were successfully applied on genomic sequences to identify lateral genetic transfers in bacterias (Cong et al. 2016), to classify genes as cancerous or not (Özcan Şimşek et al. 2019), and to try to predict the number of SARS-CoV-2 cases using genomic sequences (Vilain and Aris-Brosou 2023).

While many encoding schemes exist, choosing one to represent the genomic sequences to analyze is not always trivial. Certain numerical features are better suited for certain types of tasks or types of genomic sequences (Yu et al. 2018), often pushing researchers to test different encodings or to combine them to obtain better model performances (Choong and Lee 2017; He et al. 2019; Yuan et al. 2024). Although some software exists to generate these numerical features, many are only accessible as web servers (Chen et al. 2014; Liu et al. 2016), which is impractical when working with high-throughput analyses. Alternative options implemented in Python exist, as it is usually the preferred language to build statistical and ML models due to the many library options that exist for this programming language, such as scikit-learn (Pedregosa et al. 2011), SciPy (Virtanen et al. 2020), TensorFlow (Abadi et al. 2015), or PyTorch (Paszke et al. 2019). However, the existing options developed in Python are unsuitable for analyses at larger scales for different reasons. First, some software only offers a graphical user interface (Bonidia et al. 2022), making it impossible to integrate the method directly into other Python-based pipelines. Second, other software only provides a command line interface and does not provide options to be used within other Python scripts (Muhammod et al. 2019). Finally, to our knowledge, all Python tools available for transforming genomic sequences into numerical features are programmed solely in Python, rather than using a combination of programming languages (Bonidia et al. 2022; Muhammod et al. 2019; Chen et al. 2022). As a highlevel programming language, Python trades runtime performance for flexibility and ease of use (Schofield and Hodson 2024), which potentially leads to longer encoding times when transforming many sequences. Because data preprocessing, including data transformation, has been reported to take 50 to 80 percent of data scientists’ time (Lohr 2014) and constitute 42 percent of the steps in bioinformatic pipelines (Chen and Chang 2017), streamlining preprocessing steps is crucial, especially as datasets grow in size.

To solve these shortcomings, we propose `dna-parser`, a Python library written in Rust to encode large datasets of genomic sequences. We describe a flexible implementation allowing users to integrate it into other Python scripts while providing a significant speed-up in encoding times through Rust. The library provides ten widely used numerical feature schemes from bioinformatics and natural language processing. Furthermore, we test the encoding speed of our implementation against two popular bioinformatics Python packages to encode DNA and RNA sequences, MathFeature (Bonidia et al. 2022) and IFeatureOmega (Chen et al. 2022). We also test `dna-parser` implementation against a broadly used natural language processing library, scikit-learn, which enables text data encoding and transformation, as well as building machine learning models.

## 2 Implementation

Through its implementation, `dna-parser` maintains the flexibility provided by the Python programming language and compatibility with other packages in this ecosystem while providing fast encoding of genomic sequences. This is achieved using two programming languages to implement `dna-parser`, Python and Rust. The Python side acts as a wrapper around the Rust side, allowing users to interact with the encoding functions written in Rust. Additionally, it returns encodings in a NumPy format (Harris et al. 2020) that can be directly used by other libraries such as scikit-learn, SciPy, TensorFlow, or PyTorch. Rust, a low-level compiled programming language, is used to implement the encoding functions. Using Rust provides three main advantages. First, it is a faster programming language than Python. Second, the Python Global Lock Interpreter (GIL) prevents Python from running on multiple threads simultaneously. Once the sequences to encode are transferred to the Rust side, the GIL can be released. This allows Python to perform other actions while the sequences are being encoded on the Rust side. Third, Rust enables parallelization, meaning that multiple sequences can be encoded at once on different threads. Finally, the Python and Rust sides are connected using PyO3 (PyO3 Project and Contributors .), a package that manages acquiring or releasing the GIL and that performs data type conversions between the two programming languages.

### 2.1 dna-parser workflow and functionalities

`dna-parser` was created to encode many genomic sequences into numerical descriptors. It provides the option to load sequences from FASTA files, but can also encode sequences already loaded in memory, providing more flexibility. Furthermore, the sequences can be encoded in parallel using multiple threads, resulting in additional encoding speed up if necessary. `dna-parser` offers many numerical feature schemes coming from bioinformatics such as Atomic Number (Kwan and Arniker 2009), Chaos Game representation (Jeffrey 1990), Cross encoding (inspired from complex number representation) (Kwan and Arniker 2009), DNA walk (Hewelt et al. 2019), EIIP (Nair and Sreenadhan 2006), Fickett Score (Fickett 1982), real-number (Chakravarthy et al. 2004), and Z-curve representations (Zhang and Zhang 2014). Some encodings used in the field of natural language processing but relevant for genomic sequences like the one-hot (or Voss) (Voss 1992; Kwan and Arniker 2009; Yu et al. 2018) encoding or the TF-IDF (Salton and Buckley 1988) are also implemented. The encodings are returned as a NumPy array or as a SciPy sparse matrix for the TF-IDF encoding. A complete list of encodings with examples and explanations of function parameters can be found at: https://mvila035.github.io/dna_parser/.

### 2.2 Benchmarks

To test and compare `dna-parser` against other software, we collected representative genomic sequences to encode them into numerical features. We used Genomic Bench-marks (https://github.com/ML-Bioinfo-CEITEC/genomic_benchmarks), a repository that gathers datasets for benchmarking the classification of genomic sequences (Grešová et al. 2023). From Genomic Benchmarks, we downloaded all of the available datasets. After plotting the distribution of sequence lengths, we noticed that some were abnormally short (as short as 2 bp). Thus, we removed sequences shorter than 100 bp, resulting in 877,848 sequences. The software we benchmark against assumes the sequences to encode are aligned and thus have equal length. Padding all sequences to the longest one could lead to misleading results, as encoding sequences with highly repetitive characters is more predictable by the Central Processing Unit (CPU) and may result in faster encoding times. Thus, we decided instead to truncate every sequence down to 100 bp.

Next, we loaded the sequences in memory and measured the time taken to encode all sequences for each of the descriptors implemented in `dna-parser` using a single CPU thread. Then, we tested the encoding implementation of `dna-parser` against MathFeature and IFeatureOmega. We measure the encoding time for the first 10, 50, 100, 1000, 5000, 10,000, 50,000, 100,000, 500,000, 877,848 (full dataset), 4,389,240 (five times the full dataset), and 8,778,480 (10 times the full dataset) sequences using the EIIP numerical feature scheme. To assess the effect of sequence length, we conducted a second benchmark against MathFeature and IFeatureOmega. We either truncated each sequence, or extended it by repeating the sequence until the desired length was reached. We measured encoding times for the full dataset with sequence lengths of 10, 50, 100, 300, 500, 1,000, 3,000, and 5,000 bp. Additionally, we benchmarked the TF-IDF implementation in `dna-parser` against the one implemented in the scikit-learn library, as neither MathFeature or IfeatureOmega support this encoding. The TF-IDF scheme was benchmarked using *k*-mers of size 12 and the same workflow as the EIIP encoding. Each benchmark involving `dna-parser` was first performed using only a single thread to have a direct comparison with MathFeature, IFeatureOmega, and scikit-learn, which are single-threaded programs. Then the benchmarks were repeated using 2, 4, 8, and 16 threads to gauge how multiple threads improves performance. Each measurement for each benchmark was repeated five times to obtain a more robust estimate of encoding performance. Finally, all benchmarks were conducted on the Cedar supercomputer at the Digital Research Alliance of Canada.

## 3 Results and Discussion

Before conducting any benchmark, we expected some encoding schemes to be faster than others because some merely map each bp to a value, while others are more sequential. The results presented in Table 1 support this intuition. Schemes like Atomic Number, Cross encoding, EIIP, one-hot encoding, real-number encoding, Chaos Game, DNA walk, Fickett Score, TF-IDF, and Z-Curve encode faster than DNA walk, Fickett Score, TF-IDF, and Z-Curve. Out of every encoding scheme, TF-IDF was by far the slowest, as this algorithm involves mapping *k*-mers frequency within each sequence, calculating sequence frequencies for all *k*-mers, and calculating the TF-IDF scores for each sequence.

**Table 1.** Summary of descriptors implemented in dna-parser along with the time taken to encode all sequences in the benchmark dataset. All benchmarks were run using a single thread.

| Name | Encoding Time<br>(in seconds) | Encoding<br>Dimensions | Reference |
| --- | --- | --- | --- |
| Atomic Number | 0.78 | $n \times l$ | (Kwan and Arniker 2009) |
| Chaos Game | 1.10 | $n \times l \times 2$ | (Jeffrey 1990) |
| Cross | 0.82 | $n \times l \times 2$ | (Kwan and Arniker 2009) |
| DNA Walk | 1.18 | $n \times l \times 2$ | (Hewelt et al. 2019) |
| EIIP | 0.91 | $n \times l$ | (Nair and Sreenadhan 2006) |
| Fickett Score | 1.21 | $n$ | (Fickett 1982) |
| One-hot (Voss) | 0.89 | $n \times l \times 4$ | (Yu et al. 2018) |
| Real-number | 0.83 | $n \times l$ | (Chakravarthy et al. 2004) |
| TF-IDF | 8.10 | $n \times k$ | (Salton and Buckley 1988) |
| Z-Curve | 3.09 | $n \times l \times 3$ | (Zhang and Zhang 2014) |

$n$ : number of sequences; $l$ : length of sequences (in base pair); $k$ : number of unique $k$ -mers across all sequences.

We also expected our implementation to be multiple times faster than other libraries because Rust is a lower-level compiled programming language that is much faster than Python. The results in Figure 1 a) indicate that `dna-parser` consistently outperformed other existing libraries for encoding speeds when encoding more than 25 sequences of 100 bp. Below 25 sequences, the overhead of calling Rust from Python and converting the data types between the two languages results in slightly longer encoding times. However, the time to encode 25 sequences is less than 0.25 milliseconds for all libraries; thus, the difference in encoding speed is not very critical at this scale. Above this number of sequences, `dna-parser` was on average 5.9 times faster than MathFeature (Bonidia et al. 2022) and 9.8 times faster than IFeatureOmega (Chen et al. 2022) when using a single thread.

**Fig. 1.**
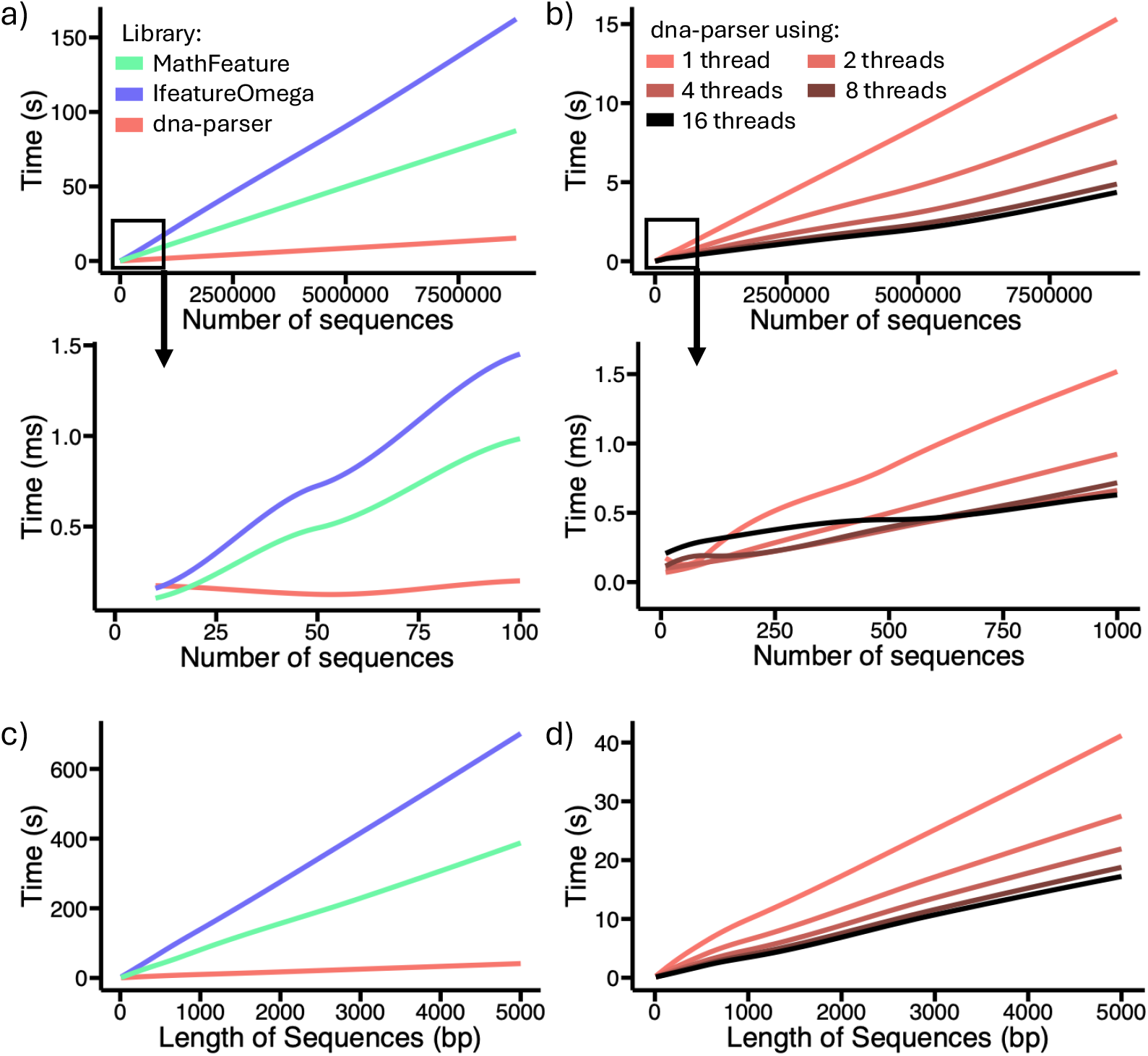
Benchmark results for the EIIP encoding scheme. **a)** Comparison of the time taken to encode sequences between `dna-parser`, MathFeature, and IFeatureOmega libraries using a single thread. The lower graphic presents a zoomed-in view of the upper, one focusing on smaller numbers of sequences to encode. **b)** Comparison of the time taken to encode sequences when using `dna-parser` with different numbers of threads. The lower graphic presents a zoomed-in view of the upper one, focusing on smaller numbers of sequences to encode. **c)** Comparison of the time taken to encode 877,848 sequences of various lengths between `dna-parser`, MathFeature, and IFeatureOmega libraries using one thread. **d)** Comparison of the time taken to encode 877,848 sequences of various lengths when using `dna-parser` with different numbers of threads.

Additionally, using multiple threads further increased the performance of `dna-parser`, as shown in Figure 1 b). When using 16 threads to encode the maximum number of sequences in our benchmark, our implementation was 20 and 37.2 times faster than MathFeature and IFeatureOmega, respectively. However, it is important to note that creating threads, passing data to them, and collecting results has a cost or overhead. Indeed, using 16 threads only leads to better performance than a single thread when encoding 200 sequences or more. This is the point at which the performance gained becomes greater than the overhead.

When varying the length of sequences to encode, we observe a similar improvement in encoding times. In Figure 1 c), for a single CPU thread, `dna-parser` was on average 7.0 times faster than MathFeature and 12.8 times faster than IFeatureOmega. Moreover, in figure Figure 1 d), when using 16 threads to encode sequences that were 5000 bp long, `dna-parser` was 22.5 and 40.7 times faster than MathFeature and IFeatureOmega, respectively.

We compared our TF-IDF implementation to the one in the scikit-learn library. When varying the number of sequences to encode, `dna-parser` was always faster than scikit-learn, even for a small number of sequences (Figure 2 a) and b)). However, we did not obtain as much speed improvement compared to the EIIP benchmark. `dna-parser` was on average 2.6 times faster than scikit-learn. This smaller speed improvement can be explained by the very efficient implementation of the TF-IDF due to the popularity of the scikit-learn library. When using eight threads and encoding the most amount of sequences, `dna-parser` was 3.4 times faster than scikit-learn. Surprisingly, using 16 threads was always slower than using only 8 for the range of sequences to encode that we benchmarked. This could be due to the short length of sequences (100 bp). One of the main parallelizable steps in the TF-IDF is to split the sequences into *k*-mers and count how many appear in each sequence. But, for short sequences, the performance gained by using multiple threads might not outweigh the cost of spawning said threads.

**Fig. 2.**
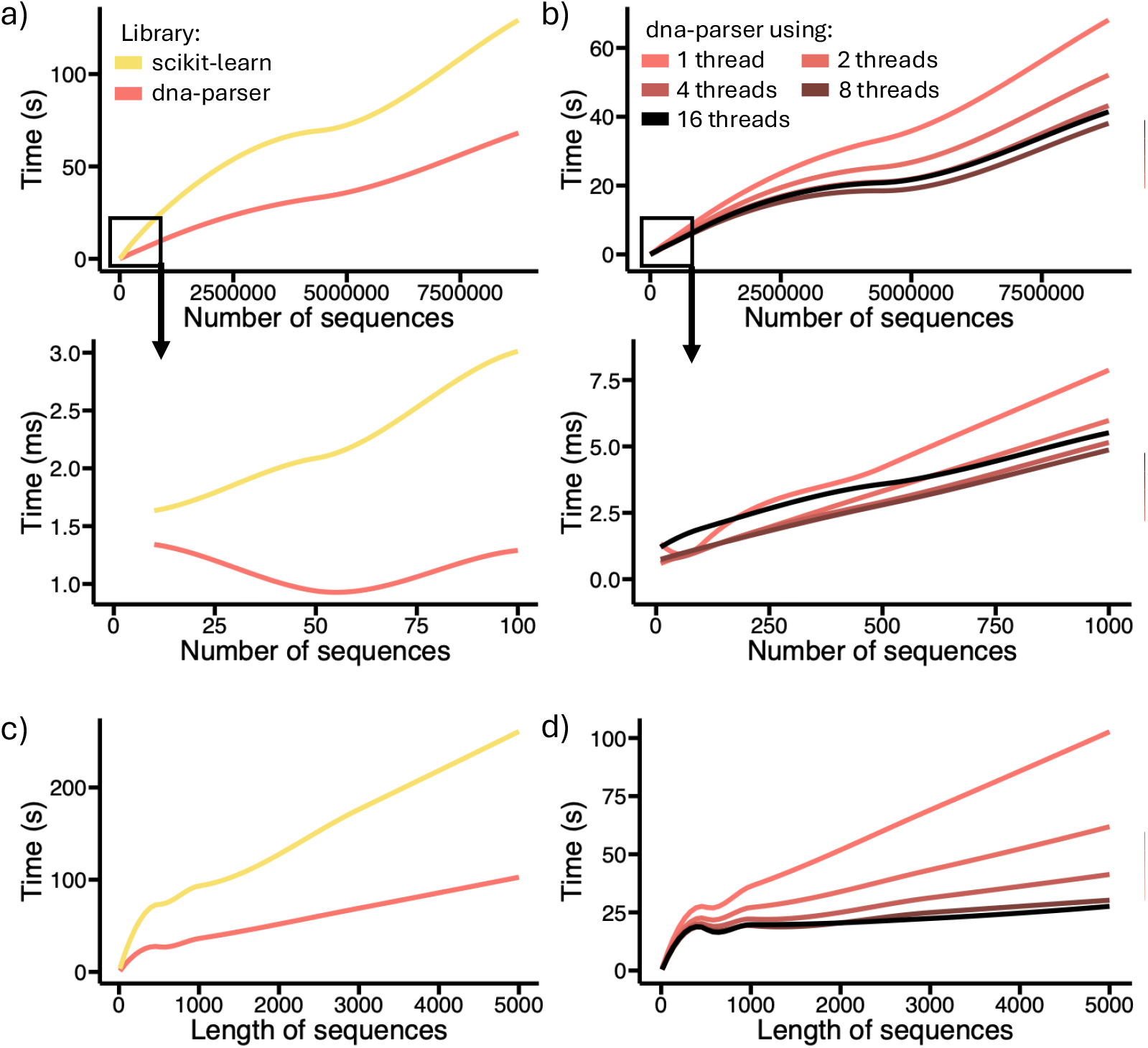
Benchmark results for the TF-IDF encoding scheme. **a)** Comparison of the time taken to encode sequences between `dna-parser` and scikit-learn libraries using one thread. The lower graphic presents a zoomed-in view of the upper, one focusing on smaller numbers of sequences to encode. **b)** Comparison of the time taken to encode sequences when using `dna-parser` with different numbers of threads. The lower graphic presents a zoomed-in view of the upper one, focusing on smaller numbers of sequences to encode. **c)** Comparison of the time taken to encode 877,848 sequences of various lengths between `dna-parser` and scikit-learn libraries using one thread. **d)** Comparison of the time taken to encode 877,848 sequences of various lengths when using `dna-parser` with different numbers of threads.

When varying the length of the sequences, we obtained similar results. In Figure 2 c), the `dna-parser` implementation using a single thread was on average 2.7 times faster than scikit-learn. Moreover, encoding longer sequences led to better performance when using multiple threads. As shown in Figure 2 d), running only eight threads was faster than using 16 until the length of sequences reached 2000 bp. After this threshold, using `dna-parser` with 16 threads led to better performance and was up to 9.4 times faster than scikit-learn when encoding sequences that were 5000 bp long. While `dna-parser` already offers a considerable speed up to encode large datasets, other strategies could be implemented to increase performances further. Single Instruction on Multiple Data (SIMD) (Flynn 2009) allows the CPU to perform the same operation simultaneously on different data. This type of parallel computing could provide an additional speedup for encoding schemes mapping each bp to a value. The Rust compiler, when using the “-C target -cpu=native” flags, can automatically detect if a machine CPU supports some SIMD instructions and can try to automatically vectorize some functions at compilation time. However, there is no guarantee that all if any function will be vectorized. Alternatively, the Rust encoding functions could be rewritten using specific SIMD instructions. Additionally, encoding functions could also be rewritten using Graphics Processing Units (GPU) instructions to take advantage of their highly parallel architecture. However, encoding sequences on a GPU entails transferring the sequences to the GPU memory and might not always result in faster encoding speed than using SIMD instructions.

Overall, we decided not to adopt these approaches as they would likely render the accessibility and distribution of our library less universal. Indeed, to create Python wheels with specific SIMD instructions, we would need to compile for each combination of Python version, exploitation system, CPU architecture, and SIMD instruction; many combinations to which we do not have access and could not compile. Furthermore, the same problem arises with GPUs as they do not all support the same instructions, and thus compiling for each type of GPU would be necessary.

## 4 Conclusion

`dna-parser` is a Python package that can be used to encode DNA and RNA sequences into numerical features faster. This package uses a combination of Rust and Python to encode sequences up to 40 times faster than other state of the art package, while retaining flexibility for the users. Being fast and flexible, `dna-parser` is an excellent candidate to be integrated in high-throughput pipelines that need to convert DNA/RNA sequences into numerical representations. Additionally, the library offers many prebuilt Wheels for Python 3.9 to 3.13 on Linux, macOS and Windows platform that can be easily installed via pip. This package will help researchers creating ML pipelines to rapidly test different encodings or to deploy their pipeline at scale. Finally, this software also showcases how Rust and Python can be an excellent combination to build efficient bioinformatics tools.

## 5 Availability and requirements

**Project name:** dna-parser

**Project home page:** https://github.com/Mvila035/dna_parser

**Operating system(s):** Windows, macOs, Linux

**Programming language:** Python, Rust

**Other requirements:** Python 3.9 or higher

**License:** MIT

**Any restrictions to use by non-academics:** None

## 6. List of abbreviation

bp: Base Pair
CPU: Central Processing Unit
EIIP: Electron-Ion Interaction Pseudopotential
GPU: Graphics Processing Unit
ML: Machine Learning
SRA: Sequence Read Archive
SIMD: Single Instruction on Multiple Data
TF-IDF: Term Frequency-Inverse Document Frequency

## 7 Declarations

### 7.1 Ethics approval and consent to participate

Not Applicable

### 7.2 Consent for publication

Not Applicable

### 7.3 Availability of data and materials

The data that support the findings of this study was acquired through the online repository Genomic Benchmarks: https://doi.org/10.1186/s12863-023-01123-8. Instructions on how to use the repository can be found at: https://github.com/ML-Bioinfo-CEITEC/genomic_benchmarks

### 7.6 Authors’ contributions

M.V. conceived the software, M.V. and S.A.-B. conceived the methodology,

M.V. conducted the formal analysis, M.V. wrote the original manuscript draft,

M.V. and S.A.-B. reviewed and edited the manuscript, S.A-B. acquired the funding, S.A.-B. supervised the project. All authors have read and agreed to the published version of the manuscript.

## 7.7 Acknowledgements

The authors thank the Digital Research Alliance of Canada for providing us with access to their high performance clusters.

